# Quantitative analysis of signaling responses during mouse primordial germ cell specification

**DOI:** 10.1101/2021.01.11.426293

**Authors:** Sophie M. Morgani, Anna-Katerina Hadjantonakis

## Abstract

During early mammalian development, the pluripotent cells of the embryo are exposed to a combination of signals that drive exit from pluripotency and germ layer differentiation. At the same time, a small population of pluripotent cells give rise to the primordial germ cells (PGCs), the precursors of the sperm and egg, which pass on heritable genetic information to the next generation. Despite the importance of PGCs, it remains unclear how they are first segregated from the soma, and if this involves distinct responses to their signaling environment. To investigate this question, we mapped BMP, MAPK and WNT signaling responses over time in PGCs and their surrounding niche *in vitro* and *in vivo* at single-cell resolution. We showed that, in the mouse embryo, early PGCs exhibit lower BMP and MAPK responses compared to neighboring extraembryonic mesoderm cells, suggesting the emergence of distinct signaling regulatory mechanisms in the germline versus soma. In contrast, PGCs and somatic cells responded comparably to WNT, indicating that this signal alone is not sufficient to promote somatic differentiation. Finally, we investigated the requirement of a BMP response for these cell fate decisions. We found that cell lines with a mutation in the BMP receptor (*Bmpr1a*^−/−^), which exhibit an impaired BMP signaling response, can efficiently generate PGC-like cells revealing that canonical BMP signaling is not cell autonomously required to direct PGC-like differentiation.

## Introduction

Primordial germ cells (PGCs) are the embryonic precursors of the sperm and egg that are required to pass on heritable genetic information to the next generation. Defects in PGC production result in infertility while transformed or incorrectly positioned PGCs may give rise to germ cell tumors [1–4]. Thus, delineating the mechanisms that control PGC formation is critical to our understanding of both development and disease.

In mouse, PGCs emerge during early development at a time when the pluripotent cells of the embryo are exposed to a myriad of signals that drive cell fate specification. These signals direct the majority of cells to adopt somatic fates [5], while a small population of only around 40 cells repress the somatic program and instead become PGCs [6–8]. Despite the importance of these cells, it is unclear how distinct germline and soma identities emerge within a common signaling environment. Unlike somatic cells, PGCs express pluripotency-associated factors, including Oct4 (Pou5f1), Sox2, Nanog, Alkaline Phosphatase, and Ssea-1 [9] and demonstrate pluripotent properties, such as the capacity to give rise to self-renewing cell lines *in vitro*, and teratomas *in vivo* [10, 11]. This has led to the hypothesis suggesting that PGCs are the last cells of the embryo to differentiate [12], and thus may not initially respond to differentiation cues. Consistent with this notion, while emerging in a region of the embryo that is exposed to high levels of Bone Morphogenetic Protein (BMP) signaling factor, PGCs do not exhibit a BMP signaling response although their immediate somatic neighbors do [13, 14]. Nevertheless, we still know almost nothing about how PGCs respond to the other biochemical signals present within their environment in the embryo and how these responses change over time.

To address this, we systematically and quantitatively analyzed the response of individual PGCs and neighboring somatic niche cells to key signals present within the embryo during PGC specification. We confirmed that PGC-like cells (PGCLCs), generated from embryonic stem cells (ESCs) *in vitro*, and PGCs *in vivo* displayed significantly lower BMP signaling responses than non-PGCs. We found that early PGCs *in vivo* also show a diminished Mitogen-Activated Protein Kinase (MAPK) response, revealing PGC-specific modes of signaling regulation for multiple pathways. In contrast, PGCs responded to WNT comparably to somatic niche cells. Therefore, PGCs are not refractory to all signals within their environment and, in this context, in the absence of robust BMP and MAPK responses, WNT signaling is not sufficient to drive somatic differentiation in cells to be assigned a PGC fate.

Finally, while PGCs are devoid of a BMP signaling response, BMP is required for PGC specification [15–20], but its role and mechanism of action remain elusive. Here, we showed that ESCs with a mutation in the BMP receptor type Ia (*Bmpr1a*) gene, which are defective in their canonical BMP signaling response, can efficiently generate PGCLCs revealing that a robust canonical BMP response is neither required transiently at earlier stages of differentiation, or indirectly via the somatic niche for early PGC differentiation.

## Methods

### 2.1 Cell culture and PGCLC *in vitro* differentiation

Cells were maintained at 37°C, at 5% CO_2_ and 90% humidity. ESC lines were routinely cultured in serum/LIF medium (Dulbecco’s modified Eagle’s medium (DMEM) (Gibco, Gaithersburg, MD) containing 0.1 mM non-essential amino-acids (NEAA), 2 mM glutamine and 1 mM sodium pyruvate, 100 U/ml Penicillin, 100 μg/ml Streptomycin (all from Life Technologies, Carlsbad, CA), 0.1 mM 2-mercaptoethanol (Sigma, St. Louis, MO), and 10% Fetal Calf Serum (FCS, F2442, Sigma) and 1000 U/ml LIF on plates coated with 0.1 % gelatin, as described [21]. The following cell lines were used in this study: E14 (129/Ola background) [22], TCF/Lef:H2B-GFP [23], *Spry4*^H2B-Venus^ [24], and *Bmpr1a*^−/−^ [25].

*In vitro* PGC-like cell (PGCLC) differentiation was performed as described [26]. Briefly, ESCs were converted to an epiblast-like (EpiLC) state by 48 hour culture in N2B27 medium containing 12 ng/ml FGF2 (233-FB-025, R&D Systems) and 20 ng/ml ACTIVIN A (120-14P, Peprotech, Rocky Hills, NJ) on dishes coated with 16.7 μg/mL fibronectin (FC010, Millipore). Following EpiLC conversion, cells were trypsinized to a single cell suspension and 10,000 cells/mL were resuspended in PGCLC medium, comprising GMEM (Gibco), 0.1 mM NEAA, 2 mM glutamine and 1 mM sodium pyruvate, 100 U/ml Penicillin, 100 μg/mL Streptomycin, 0.1 mM 2-mercaptoethanol, 1000 U/ml LIF, 15 % Knockout serum replacement, with 500 ng/ml BMP4, 500 ng/ml BMP8a, 100 ng/ml SCF, and 50 ng/ml EGF (all from R&D Systems). Samples were collected for analysis at day 0 (EpiLC state), 2, 4 and 6 of differentiation.

### 2.2 Flow cytometry

Between 8-12 PGCLC aggregates per cell line/condition were pooled and then dissociated by incubation in TrpLE™ Select Enzyme (Thermo Fisher Scientific) at 37°C for approximately 2 minutes. Following vigorous pipetting to form a single-cell suspension, the enzyme was neutralized with an equal volume of PGCLC medium without cytokines added. Cells were pelleted by centrifugation and then resuspended in 100 μL FACs buffer (PBS with 10 % FCS) with PE-conjugated anti-CD61 (RRID:AB_313084, Biolegend, 104307, 1:200) and Alexa Fluor 647-conjugated anti-SSEA1 (RRID:AB_1210551, Thermo Fisher Scientific, 51-8813-73, 1:50) for 15 min on ice. Cells were then washed in 1 mL FACS buffer and resuspended in 200 μL FACS buffer containing 5◻μg/ml Hoechst. Samples were analyzed using a BD LSR Fortessa™. Flow cytometry analysis was performed using FlowJo software (BD Biosciences). Cells were first separated from debris and cell doublets removed by gating on forward (FSC) and side scatter (SSC). Subsequently, dead cells were identified based on strong Hoechst staining and were excluded from further analysis. Gating for CD61, SSEA-1 positive cells was based on unstained wildtype E14 ESCs.

### 2.3 Mouse lines

Mice were housed under a 12◻hr light-dark cycle in a pathogen-free room in the designated MSKCC facilities. For this study we used outbred CD1 animals maintained in accordance with the guidelines of the Memorial Sloan Kettering Cancer Center (MSKCC) Institutional Animal Care and Use Committee (IACUC). Natural mating was set up in the evening and mice were checked for copulation plugs the next morning. The date of vaginal plug was estimated as E0.5. For analysis of post-implantation stages of development, embryos were isolated from deciduae and Reichert’s membrane removed by microdissection before further processing.

### 2.4 Immunostaining

Cell lines were immunostained as previously described [21]. Post-implantation embryos were washed in phosphate-buffered saline (PBS), then fixed in 4 % paraformaldehyde (PFA) for 15 min at room temperature (RT). Embryos were washed in PBS plus 0.1 % Triton-X (PBST-T) followed by permeabilization for 30 min in PBS with 0.5 % Triton-X. Embryos were then washed in PBS-T and blocked overnight at 4 °C in PBS-T with 1 % bovine serum albumin (BSA, Sigma) and 5 % donkey serum (Sigma). The following day, embryos were transferred to the primary antibody solution (PBS-T with appropriate concentration of antibody) and incubated overnight at 4 °C. The following day, embryos were washed 3 × 10 min in PBS-T and transferred to blocking solution at RT for a minimum of 5 hr. Embryos were transferred to secondary antibody solution (PBS-T with 1:500 dilution of appropriate secondary conjugated antibody and 5◻μg/ml Hoechst) overnight at 4 °C. Embryos were washed 3 × 10 min in PBS-T.

The following primary antibodies were used in this study: AP2_γ_ (RRID:AB_667770, Santa Cruz, sc-12762, 1:100), phosphorylated SMAD1/5/9 (a gift from Dr. Edward Laufer, University of Utah School of Medicine), Sox2 (RRID:AB_11219471, Thermo Fisher Scientific, 14-9811-82, 1:200).

### 2.5 Cryosectioning

Following wholemount immunostaining and imaging, embryos were oriented as desired and embedded in Tissue-Tek^®^ OCT (Sakura Finetek, Japan). Samples were frozen on dry ice for approximately 30 min and then maintained for short periods at −80◻°C followed by cryosectioning using a Leica CM3050S cryostat. Transverse cryosections of 10◻μm thickness were cut with a Leica CM3050S cryostat and mounted on Colorfrost Plus^®^ microscope slides (Fisher Scientific) using Fluoromount G (RRID:SCR_015961, Southern Biotech, Birmingham, AL). Cryosections were then imaged using a confocal microscope as described.

### 2.6 Quantitative image analysis

Embryos were imaged on a Zeiss LSM880 laser scanning confocal microscope. Confocal z stacks of cells or embryo cryosections were generated. Raw data was then processed in ImageJ open source image processing software (Version: 2.0.0-rc-49/1.51d). Individual PGCLCs, identified by AP2γ expression, PGCs identified by SOX2 expression, or their surrounding AP2γ- SOX2- niche cells were randomly chosen and, using Fiji (ImageJ) software, selected by manually drawing a boundary around the nucleus. The mean fluorescence intensity of pSMAD1/5/9 immunostaining, *Spry4^H2B-Venus^*, or TCF/Lef:H2B-GFP reporter expression was then measured in arbitrary units. Fluorescence decay along the z-axis was corrected for each channel and sample by fitting a linear regression model to the logarithm of fluorescence values as a function of the z-value, and correcting the models’ slopes using an empirical Bayes approach, as previously described [27]. For all quantification, statistical analysis of significance was assessed using a One-way ANOVA followed by unpaired *t*-tests to compare particular groups (GraphPad Prism, GraphPad Software, Inc., Version 7.0a). For analysis performed on embryos, all PGCs were selected from 3 different cryosections through the allantois of 3 distinct embryos. Fluorescence values were then calculated relative to the average mean fluorescence of non-neighboring (‘Other’) AP2γ- SOX2- niche cells within each individual section in order to normalize for differences in immunostaining that may arise due to differences in permeability within different embryonic regions or different stages of development. Statistics were carried out on average fluorescence levels per embryo, rather than on a per cell basis.

## Results and discussion

### 3.1 Quantitative analysis of signaling responses during mouse PGCLC specification

Functional PGC-like cells (PGCLCs) can be generated *in vitro* from mouse embryonic stem cells (ESCs). First ESCs are converted to an epiblast-like cell (EpiLC) state, comparable to the pluripotent embryonic cells before germ layer differentiation. Subsequently, EpiLCs are aggregated in suspension culture and exposed to a combination of signals, mimicking those present in the embryo, that promote PGC specification, survival, and proliferation (Fig. 1A) [26]. Using this protocol, we successfully generated PGCLCs, identified by the coexpression of SOX2 and AP2γ (Fig. 1B), and the cell surface markers SSEA-1 and CD61 (Fig. 1C, D) [28]. PGCLC aggregates displayed widespread SOX2 expression while AP2γ was expressed in only a subset of cells, suggesting that the rate of PGCLC specification was variable across individual cells or that a mixture of cell fates were formed (Fig. 1B). Thus, we considered PGCLCs as cells that coexpressed SOX2 and AP2γ in our downstream analyses. Using this cell culture system, we then analyzed signaling responses in individual PGCLCs and surrounding non-PGCLCs.

**Figure 1.**
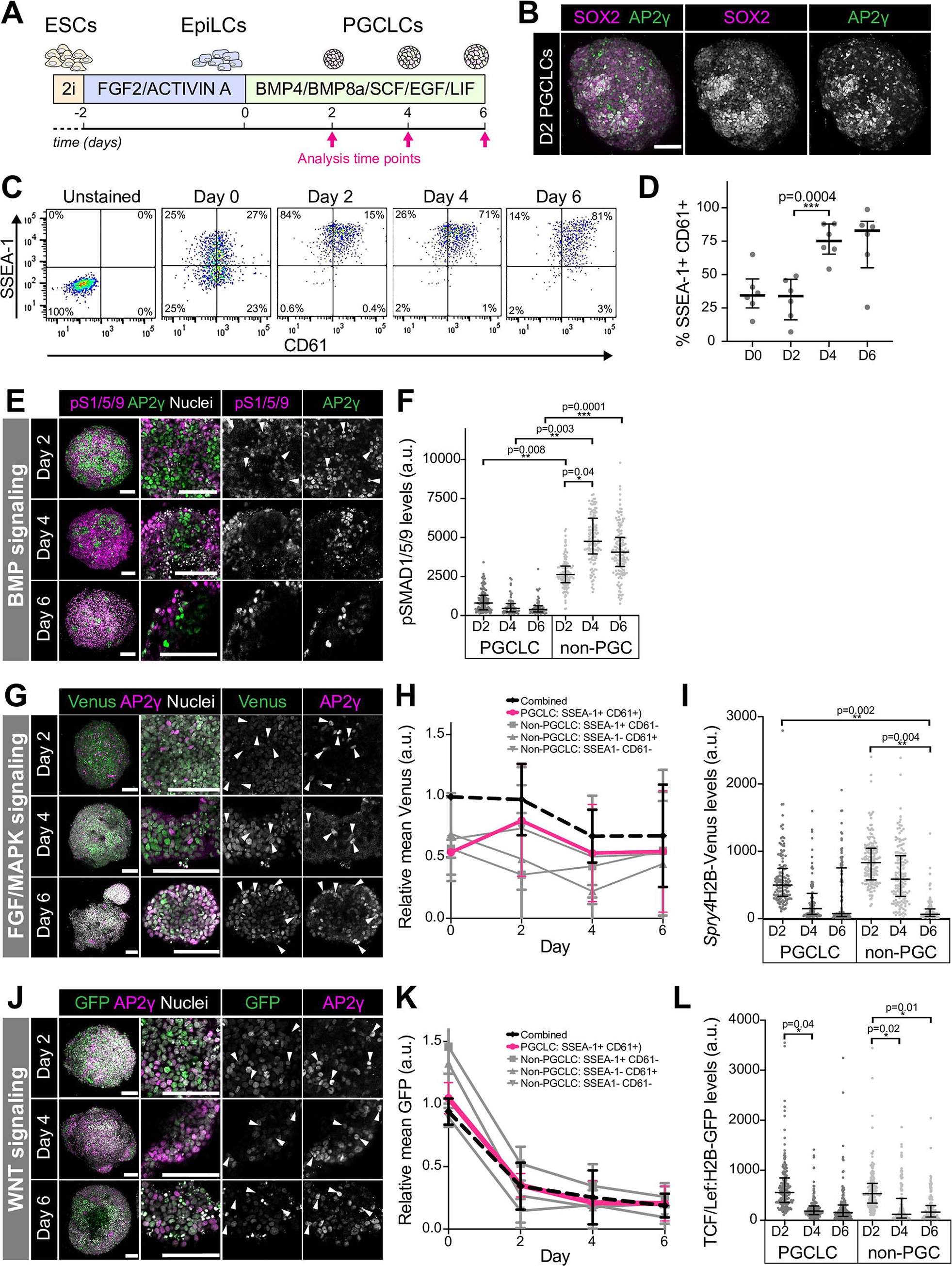
Quantitative analysis of signaling responses during PGCLC differentiation. **A.** Schematic diagram depicting the PGCLC differentiation protocol as previously described [26]. **B.** Confocal maximum intensity projection of an aggregate of cells at Day 2 (D2) of PGCLC differentiation. Scale bars, 100 μm. **C.** Representative flow cytometry data of embryonic stem cells during PGCLC differentiation. SSEA-1 and CD61 double positive cells mark PGCLCs. **D.** Percentage of SSEA-1+ CD61+ PGCLCs over time during PGCLC differentiation. Each point represents an independent experiment (n = 6) performed with 4 distinct cell lines. Data represented as median and interquartile range. **E, G, J.** Confocal maximum intensity projections of PGCLC aggregates at day 2, 4, and 6 of differentiation. Scale bars, 100 μm. **E.** Aggregates were immunostained for and AP2γ to mark PGCLCs and phosphorylated SMAD1/5/9 (pS1/5/9), a readout of the BMP signaling response. **G.** PGLC differentiation of *Spry4*^H2BVenus^ reporter embryonic stem cell lines, that act as a read out of FGF/MAPK signaling activity. **J.** PGLC differentiation of TCF/Lef:H2B-GFP reporter embryonic stem cell lines, which act as a read out of WNT signaling activity. **F, I, L.** Quantitative immunofluorescence analysis of signaling responses, measured in arbitrary units (a.u.), in PGCLCs (AP2γ+) and non-PGCLCs (AP2γ-) in 3 distinct cell aggregates per time point per cell line. Each point represents a single cell. Data shown as median and interquartile range. Statistical analysis of significance was assessed on log-normalized data using Student’s *t*-test, performed on the average fluorescence level in each aggregate. **H, K.** Relative *Spry4*^H2BVenus^ (H) and TCF/Lef:H2B-GFP (K) fluorescence levels in arbitrary units (a.u.) analyzed by flow cytometry in SSEA-1+ CD61+ PGCLCs, and SSEA-1-CD61+, SSEA-1+ CD61-and SSEA-1 CD61-non-PGCLC populations. Data represented as mean and standard deviation and shown relative to the mean fluorescence across all populations at day 0 of differentiation, n = 3 independent experiments.

BMP signaling plays a critical role in PGC specification. Mutations in the genes encoding *Bmp4*, *Bmp8*, and *Bmp2*, as well as the downstream effectors that mediate the BMP signaling response, *Smad1* and *Smad5*, result in a loss or significant reduction in PGC number [15–20]. Nevertheless, neither PGCLCs *in vitro* nor PGCs *in vivo* exhibit a canonical BMP signaling response [13, 14], demonstrated by the absence of nuclear-localized phosphorylated SMAD1/5/9 (pSMAD1/5/9, SMAD9 is also known as SMAD8). However, BMP responses have not been systematically and quantitatively analyzed at single-cell resolution and therefore it is unclear whether a fraction of PGCs do respond or if a transient response may occur. To investigate this, we measured pSMAD1/5/9 levels in individual nuclei within PGCLC aggregates at days 2, 4, and 6 of differentiation. We observed that AP2γ+ PGCLCs displayed significantly lower levels of nuclear pSMAD1/5/9 than AP2γ negative (AP2γ-) non-PGCLCs (Fig. 1E, F). Indeed, we did not identify any PGCLCs with nuclear-localized pSMAD1/5/9 (Fig. 1E, F). Furthermore, while the BMP signaling response increased in AP2γ- non-PGCLCs over time, it remained low in PGCLCs (Fig. 1F). Thus, at this resolution, we observed no evidence for a subset of BMP-responsive PGCLCs.

We then asked whether PGCLCs also lack responses to other critical signals present within the mouse embryo at this time. FGF is expressed within the posterior of the embryo at the time of PGC specification and is necessary for somatic germ layer specification, gastrulation EMT and concomitant cell migration [29–31]. Additionally, FGF and EGF, which both activate the MAPK pathway, are provided exogenously during PGCLC differentiation (Fig. 1A). In order to analyze the MAPK signaling response, we used a *Spry4*^H2BVenus^ ESC line, which harbors a fluorescent reporter in the endogenous locus of *Sprouty4* (*Spry4*), an early target of the pathway [24]. We observed widespread Venus expression throughout PGCLC aggregates at all stages of differentiation (Fig. 2G). We then performed flow cytometry and quantitative immunofluorescence to determine how this response changed over time. In contrast to the gradually increasing BMP response in non-PGCLCs, there was a reduction in the MAPK response over time (Fig. 1H, I). Quantitative immunofluorescence revealed no significant difference in the MAPK signaling response in PGCLCs and non-PGCLCs (Fig. 1I). At each stage of differentiation, Venus levels were lower, although not significantly, in AP2γ+ vs. AP2γ-cells (Fig. 1I).

**Figure 2.**
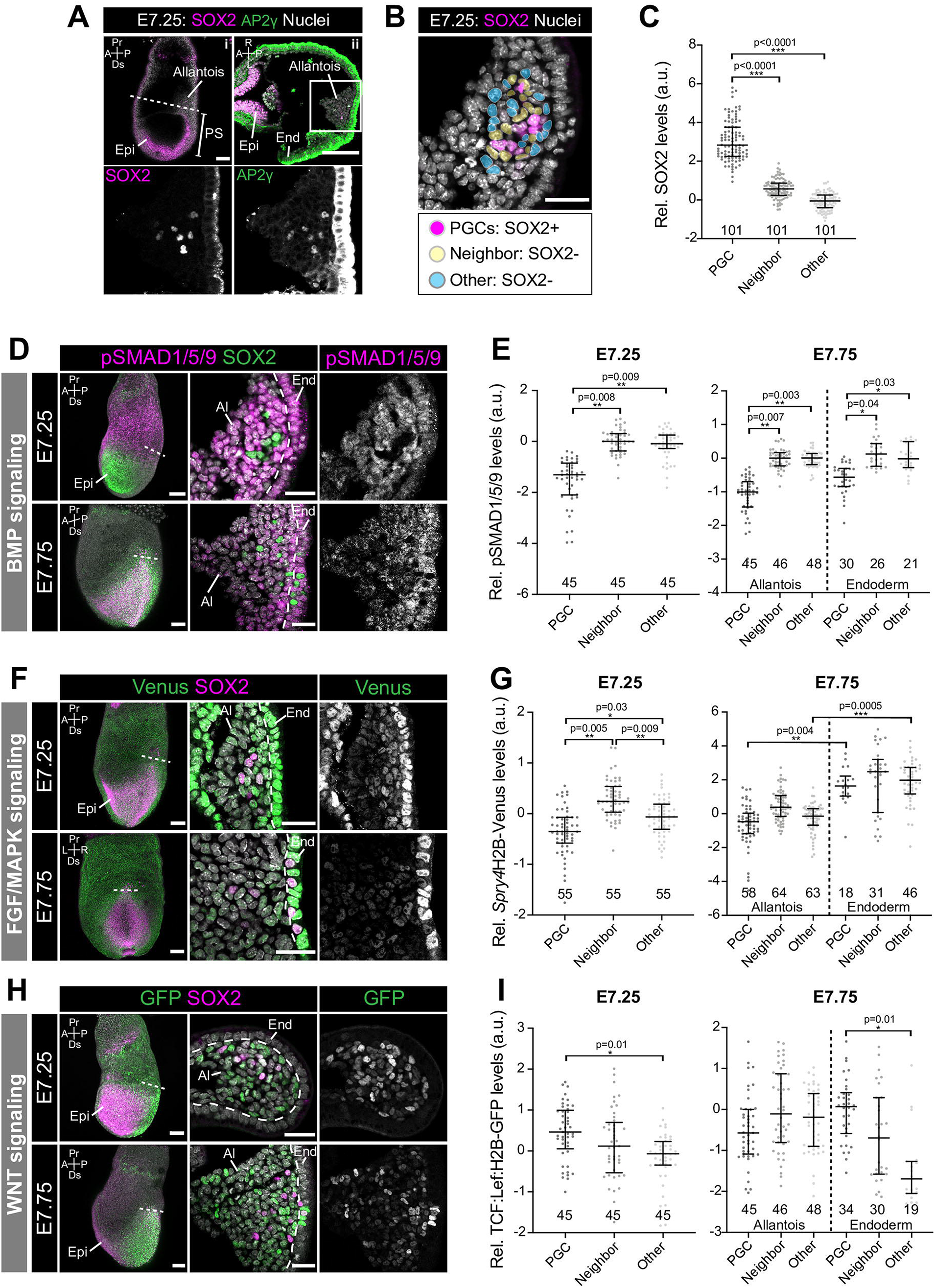
Quantitative analysis of signaling responses during PGC specification *in vivo*. **A.** (i) Sagittal confocal optical section of an immunostained embryonic day (E) 7.25 embryo. Scale bar, 100 μm. Dashed line indicates the plane of transverse section shown in adjacent panel. (ii) Confocal optical section of a transverse cryosection through the allantois of an E7.25 embryo. Scale bar, 25 μm. Box demarcates the region shown in higher magnification in lower panels. AP2γ immunostaining exhibits high levels of non-specific background staining within the endoderm. **B.** For quantitative analysis of signaling responses, cells adjacent to PGCs (pseudocolored in yellow) were categorized as PGC ‘Neighbors’ and non-adjacent cells within the allantois (pseudocolored in blue) were categorized as ‘Other’. **C.** Quantification of levels of SOX2 in arbitrary units (a.u.) PGCs, PGC Neighbors and Other cells within the allantois of E7.25 embryos. SOX2+ immunostaining was used to define the PGC population. Statistical analysis of significance was assessed using Student’s *t*-test and performed on the average fluorescence level in each embryo (n = 3 embryos, number of individual cells shown on graph). Each point represents a single cell. Data shown relative to the average mean fluorescence in ‘Other’, non-PGCs and represented as the median and interquartile range. **D, F, H.** Sagittal confocal maximum intensity projections (left panels, scale bars, 100 μm) and confocal optical sections of transverse cryosection through the allantois of E7.25 and E7.75 embryos (scale bars, 25 μm). Dashed line approximately demarcates the boundary between the allantois and the endoderm. **D.** Embryos were immunostained for phosphorylated SMAD1/5/9 as a readout of BMP signaling activity. **F.** Transgenic *Spry4*^H2BVenus^ reporter embryos were used to read out FGF/MAPK signaling activity. **H.** TCF/Lef:H2B-GFP reporter embryos were used to read out WNT signaling activity. **E, G, I.** Quantification of levels of nuclear SMAD1/5/9, *Spry4*^H2BVenus^, and TCF/Lef:H2B-GFP expression in arbitrary units (a.u.) in PGCs, PGC Neighbors and Other cells within the allantois of E7.25 and E7.75 embryos. In E7.25 embryos, all PGCs were within the allantois. In E7.75 embryos, a fraction of PGCs had also begun to migrate along the hindgut endoderm hence we separately investigated signaling responses in PGCs within the allantois and within the endoderm for this analysis. Statistical analysis of significance was assessed using Student’s *t*-test and performed on the average fluorescence level in each embryo (n = 3 embryos, number of individual cells shown on graph). Each point represents a single cell. Data shown relative to the average mean fluorescence in ‘Other’, non-PGCs and represented as the median and interquartile range. Pr, proximal; Ds, distal; A, anterior; P, posterior; L, left; R, right; Epi, epiblast; PS, primitive streak; End, endoderm.

WNT signaling is required for both somatic [32–35] and germ cell [36, 37] fate specification. WNT drives the initial exit from pluripotency but a subset of its targets must subsequently be repressed in PGC-fated cells to prevent somatic differentiation [37]. Here we used a TCF/Lef:H2B-GFP reporter ESC line, that contains multimerized binding sides for the T cell-specific transcription factor/lymphoid enhancer-binding factor 1 (TCF/Lef) family of transcription factor coactivators, which mediate the WNT signaling response [23]. Although recombinant WNT is not added exogenously to the PGCLC differentiation medium, TCF/Lef:H2B-GFP was expressed heterogeneously throughout cell aggregates (Fig. 1J), signifying that endogenous WNT ligands were present. However, there was no difference in the WNT response in PGCLCs compared to non-PGCLCs revealing that PGCLCs are not refractory to all differentiation-inducing signals. The WNT response decreased during PGCLC differentiation (Fig. 1K, L).

Thus, initially PGCLCs show a reduced BMP signaling response and as differentiation proceeds, PGCLCs and non-PGCLCs also reduce their MAPK and WNT signaling responses.

### 3.2 Quantitative analysis of signaling responses during PGC specification *in vivo*

The combination, dynamics, and dose of factors provided during PGCLC differentiation *in vitro*, may not precisely recapitulate the dynamic signaling environment within the embryo. Moreover, the majority of AP2γ- non-PGCLCs also expressed SOX2 (Fig. 1B), suggesting that they represent a pluripotent EpiLC state or earlier state of PGCLC differentiation, and thus do not mirror the *in vivo* PGC niche at the posterior of the embryo that comprises extraembryonic mesoderm. Therefore, we also sought to investigate signaling responses in PGCs and their niche in the embryo. We isolated and analyzed embryos at embryonic day (E) 7.25, when SOX2+ AP2γ+ PGCs first emerge within a posteriorly-localized extraembryonic structure known as the allantois (Fig. 2A) [38], and at E7.75, when PGCs begin to exit the allantois and migrate anteriorly along the hindgut endoderm toward their eventual destination in the gonads. In contrast to PGCLC aggregates, where only a subset of SOX2+ cells expressed AP2γ, *in vivo* SOX2 and AP2γ expression fully overlapped (Fig. 2A). However, as AP2γ immunofluorescence resulted in high levels of non-specific background staining in the endoderm on the embryo’s surface (Fig. 2A), we used SOX2 to accurately identify PGCs. We isolated wildtype embryos, which we immunostained for pSMAD1/5/9, as well as *Spry4^H2B-Venus^*, and TCF/Lef:H2B-GFP reporter embryos and measured signaling responses in individual SOX2+ PGCs, and SOX2-non-PGCs that were either adjacent to PGCs (categorized as ‘Neighbors’), or non-adjacent (categorized as ‘Other’) in transverse cryosections of the allantois (Fig. 2A, B, C). As in PGCLCs, PGCs at E7.25 and E7.75 showed significantly lower levels of nuclear-localized pSMAD1/5/9 than both neighboring and non-neighboring SOX2- cells within the surrounding somatic niche (Fig. 2D, E).

Cells within the allantois showed widespread *Spry4*^H2BVenus^ expression (Fig. 2F). However, at E7.25, PGCs displayed a significantly reduced MAPK response compared to non-PGC neighbors and non-neighboring SOX2-cells (Fig. 2F, G). By E7.75, this difference was no longer significant (Fig. 2G). PGCs within the hindgut endoderm displayed a higher MAPK response than their PGC counterparts within the allantois (Fig. 2G). Moreover, the MAPK response was higher in endoderm relative to extraembryonic mesoderm (Fig. 2G). Therefore, as PGCs migrate towards the gonads, they enter an environment of higher MAPK signaling activity.

During PGCLC differentiation there was no difference in the WNT response in PGCLCs vs. non-PGCLCs (Fig. 1L). In contrast, at E7.25 *in vivo*, PGCs expressed higher levels of TCF/Lef:H2B-GFP than non-adjacent extraembryonic mesoderm cells (Fig. 2H, I), likely a result of the distinct nature of the *in vitro* and *in vivo* niches. By E7.75, there was no difference in the WNT response between PGCs and their neighbors. However, migrating PGCs exhibited a stronger WNT response than non-adjacent endoderm. Thus, both *in vitro* and *in vivo*, PGCs respond to WNT.

### 3.3 BMP signaling response is not required for PGCLC specification

While BMP is required for PGC specification [15–20], and BMP4 and BMP8a (500 ng/UL) are exogenously provided during PGCLC differentiation [26], we and others observed that neither PGCLCs or PGCs exhibit nuclear-localized pSMAD1/5/9 (Fig. 1E, F, 2C, D) [13, 14]. Thus, either a transient BMP response is required for PGC specification that we do not capture at this temporal resolution, or BMP is required indirectly by PGCs. To distinguish between these possibilities, we performed PGCLC differentiation of *Bmpr1a*^−/−^ ESCs [25]. *Bmpr1a* is the most broadly and highly expressed BMP receptor within the pluripotent epiblast during PGC specification [39] and *Bmpr1a*^−/−^ embryos exhibit little or no nuclear pSMAD1/5/9 [40]. In keeping with this, and as previously observed [25], *Bmpr1a*^−/−^ ESCs did not display nuclear-localized pSMAD1/5/9 under standard serum/LIF culture conditions, although this was observed in wildtype *Bmpr1a*^+/+^ ESCs (Fig. 3A), or when treated with BMP4 for 2 hours (Fig. 3B). Comparable observations were made with *Bmpr1a*^−/−^ EpiLCs (Fig. 3C). We then exposed *Bmpr1a*^−/−^ EpiLCs to PGCLC induction medium and showed that, likewise, *Bmpr1a*^−/−^ cell aggregates do not exhibit nuclear-localized pSMAD1/5/9 under these conditions (Fig. 3D, E).

**Figure 3.**
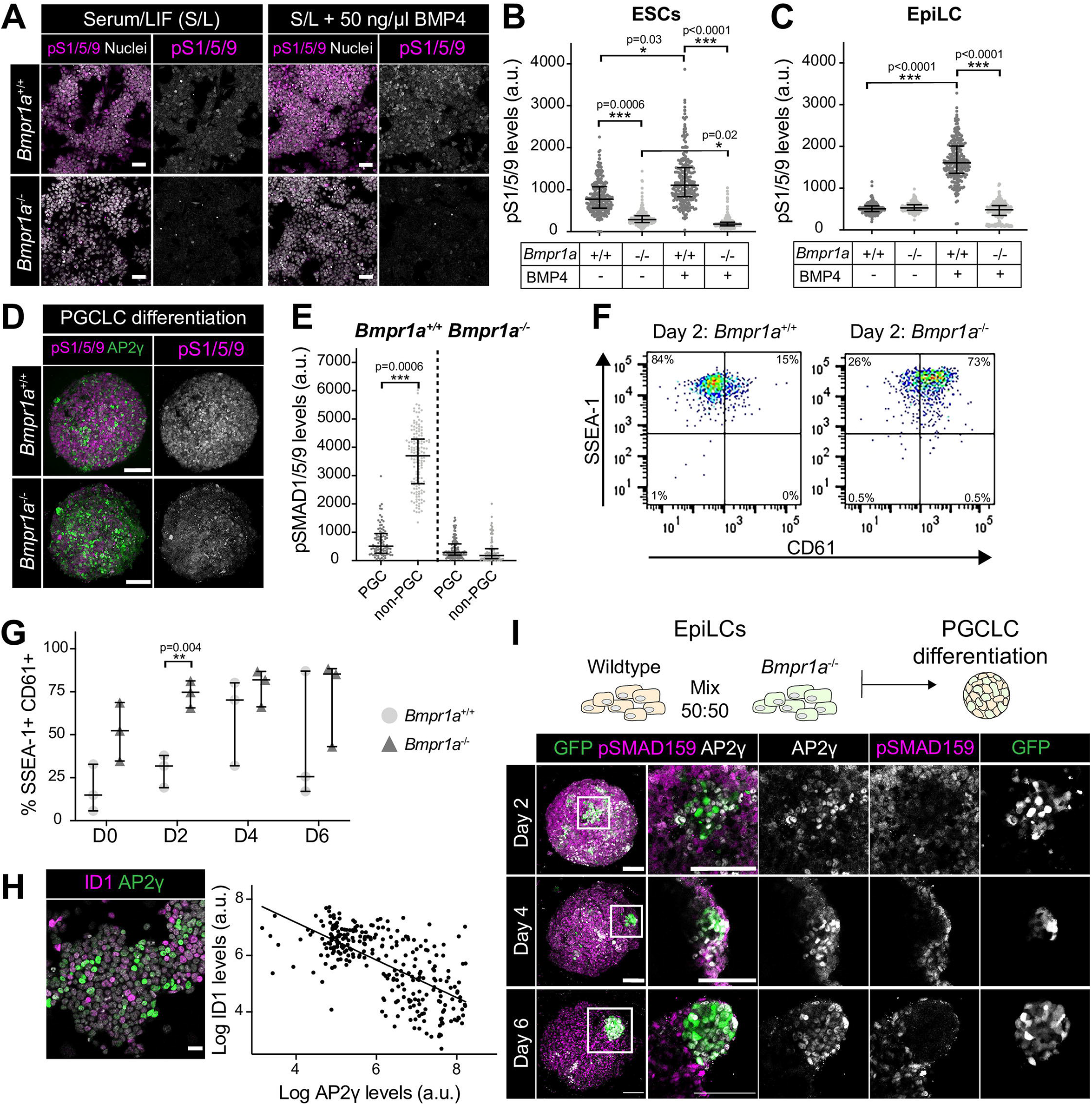
Cell autonomous BMP signaling response is not necessary for PGCLC fate. **A.** Confocal optical sections of wildtype (*Bmpr1a*^+/+^) and *Bmpr1a*^−/−^ embryonic stem cells (ESCs) immunostained for pSMAD1/5/9 (pS1/5/9) after culture under standard or after 2 hours treatment with 50 ng/ml BMP4. **B, C.** Quantification of pSMAD1/5/9 levels in wildtype and *Bmpr1a*^−/−^ embryonic stem cells ESCs (from 5 distinct fields of view) and EpiLCs (from 5 distinct fields of view). Each point represents a single cell. Data represented as median and interquartile range. Statistical analysis of significance was assessed using Student’s *t*-test, performed on the average fluorescence level in each field. n=2 experimental replicates. **D.** Confocal maximum intensity projection of wildtype and *Bmpr1a*^−/−^ cell aggregates at Day 2 (D2) of PGCLC differentiation. Scale bars, 100 μm. **E.** Quantification of pSMAD1/5/9 levels in wildtype and *Bmpr1a*^−/−^ PGCLC aggregates. Each point represents a single cell. Data represented as median and interquartile range. Statistical analysis of significance was assessed using Student’s *t*-test and performed on the average fluorescence level in each aggregate (n = 3 aggregates). **F.** Flow cytometry analysis of wildtype and *Bmpr1a*^−/−^ cell aggregates at Day 2 of PGCLC differentiation. SSEA-1+ CD61+ cells represent PGCLCs. **G.** Percentage of SSEA-1+ CD61+ PGCLCs during wildtype and *Bmpr1a*^−/−^ PGCLC differentiation. Each point represents an independent experiment (n = 3). Data represented as median and interquartile range. **H.** Confocal optical section of ESCs, cultured in serum and LIF, immunostained for the BMP pathway target, ID1 and the PGC marker AP2γ (left panel). Scale bar, 25 μm. Right panel shows quantification of ID1 and AP2γ levels in arbitrary units (a.u.) in individual cells. Quantification performed on images from 5 randomly selected regions. Each point represents a single cell. Linear regression and correlation coefficient analysis were performed and were statistically significant (p<0.0001). Correlation coefficient is shown on graph. **I.** Wildtype and *Bmpr1a*^−/−^ EpiLCs were mixed together in equal ratios to form PGCLC aggregates. *Bmpr1a*^−/−^ cells were labelled with a constitutive GFP lineage marker. Images show Confocal maximum intensity projections of PGCLC aggregates at day 2, 4, and 6 of differentiation. Scale bars, 100 μm.

Despite this, we observed the specification of cells that expressed AP2γ, as well as SSEA-1 and CD61 (Fig. 3D, F, G). Notably, *Bmpr1a*^−/−^ EpiLCs showed a slightly higher percentage of SSEA-1+ CD61+ cells than wildtype EpiLCs prior to exposure to PGCLC medium, and accordingly they displayed an earlier peak in this population during differentiation (Fig. 3G). This finding suggested that cells with a low BMP response could be predisposed towards a PGCLC fate. Consistent with this notion, we also noted an inverse correlation between the expression of the BMP pathway target Inhibitor of differentiation 1 (ID1) and the PGC marker AP2γ in wildtype ESCs (Fig. 3H). To investigate this further, we then mixed equal proportions of *Bmpr1a*^+/+^ and *Bmpr1a*^−/−^ EpiLCs, to form chimeric PGCLC aggregates (Fig. 3I). *Bmpr1a*^−/−^ ESCs were lineage labelled with a constitutive GFP reporter that enabled tracking of their eventual fate. In these experiments, we observed variable proportions of GFP+ cells within the resulting aggregates (data not shown). In aggregates with a low percentage of GFP+ cells, it was predominantly *Bmpr1a*^−/−^ cells that gave rise to and were immediately adjacent to AP2γ+ PGCLCs (Fig. 3I). Together these data indicate that a canonical BMP response is not required cell autonomously for PGCLC differentiation.

## Discussion

While many of the signals that direct PGC specification have been elucidated [14, 36, 41], little is known about how individual PGCs and their niche respond to these signals and whether their role is cell-autonomous or non-cell-autonomous. BMP is required for PGC development [15–20]. While BMP8b and BMP2 act non-cell autonomously to restrict the development of the anterior visceral endoderm, a source of PGC inhibitory signals, and to specify the hindgut required for PGC migration respectively [20, 36], the role of BMP4 is still unclear. Our and other observations [13, 14] that PGCs do not exhibit a canonical BMP signaling response, suggest that BMP4 likewise acts non-cell-autonomously or that low-levels of BMP signaling activity, not detectable following antibody staining, could be sufficient for PGC specification. This is supported by our finding that BMP signaling defective (*Bmpr1a*^−/−^) ESCs efficiently give rise to PGCLCs. However, as *Bmpr1a*^−/−^ PGCLC differentiation occurred in the absence of wildtype cells, the requirement for BMP is not via BMP-responsive cells within the niche and may instead be through non-canonical SMAD-independent downstream pathways [42, 43]. Alternatively, as perturbation of BMP signaling *in vivo* causes the epiblast to prematurely adopt a neural identity [25], it may be required to initially maintain the epiblast in a PGC competent state rather than directly regulating PGC specification. This role may not be masked *in vitro* as ESCs are forcibly maintained in a self-renewing state using 2i small molecule inhibitors.

MAPK inhibition promotes PGC differentiation *in vitro* [44]. Additionally, treating isolated PGCs with FGF reprograms them to a state of pluripotency [45]. Thus, FGF/MAPK signaling is associated with a block to the formation, or destabilization, of a PGC identity. In keeping with this, we observed that early PGCs, at E7.25, showed a lower MAPK response than somatic cells. Although we observed low-level *Spry4*^H2BVenus^ expression within PGCs and PGCLCs, indicating that MAPK signaling may not be entirely shut down, this could also represent perdurance of the Venus reporter. Therefore, future studies using dynamic ERK biosensors [46, 47], combined with time-lapse imaging [48], will uncover precise signaling dynamics. We also observed that the MAPK response was elevated in PGCs within the hindgut endoderm, consistent with studies showing that FGF plays a role in germ cell migration [45, 49]. This is at odds with reports that migrating PGCs are devoid of phosphorylated ERK, a component of the MAPK pathway [8]. Therefore, *Spry4* expression within the endoderm may also be regulated by additional signaling inputs, such as WNT [50].

PGCs are specified in a signaling environment that instructs the majority of cells to adopt a somatic non-PGC identity. One way that they might maintain their unique identity is via regulatory mechanisms that prevent them from detecting or responding to these signals. Nevertheless, while PGCs displayed a reduced BMP and MAPK response, they did respond to WNT. Hence, in the absence of robust BMP and MAPK responses, WNT is not sufficient to drive somatic differentiation. Previous studies reported that, after an initial period of WNT signaling, WNT targets must be suppressed to mediate PGC differentiation [37]. Consistent with this, we observed a rapid decrease in the WNT signaling response over time during PGCLC differentiation. However, PGCs and somatic cells showed comparable levels of TCF/Lef:H2B-GFP expression suggesting that there is not a global PGC-specific reduction in the signaling response. Thus, this regulation likely occurs via locus-specific mechanisms [37]. Of cells within the allantois, PGCs exhibit the strongest WNT response, followed by their immediate somatic neighbors, while non-neighboring somatic cells were the least responsive. Therefore, PGCs could perhaps act as a source of WNT that activates autocrine and paracrine signaling in adjacent, but not more distant cells.

Here we have shown that PGC-specific signaling responses exist for a number of different pathways. However, while we observed significant differences in MAPK and WNT signaling responses in PGCs vs. somatic cells within the allantois, these were not evident in PGCLC aggregates. This presumably reflects the difference between the PGC niche in the embryo and during stem cell differentiation, highlighting the importance of side by side comparisons of *in vitro* and *in vivo* developmental mechanisms. Furthermore, the important question remains as to how these distinct PGCs and soma responses are regulated. Current single-cell transcriptomic studies of mouse embryos contain only a small number of PGCs with no spatial information, thereby prohibiting clear conclusions on the relative expression of signaling pathway components within these cells versus their immediate neighbors and cells within their further proximity. Future PGC-enriched single-cell spatial transcriptomic studies may shed light on this. However, as signaling responses are largely regulated at a post-transcriptional level, advances in single-cell proteomic techniques or the use of quantitative time and space resolved reporters as dynamic signaling readouts may be necessary to fully answer these open questions.

## Acknowledgements

We thank members of the Hadjantonakis lab for critical discussions and comments on the manuscript. We also thank members of MSKCC’s Flow Cytometry Core facility, funded by the NCI Cancer Center Support Grant (CCSG, P30 CA08748). Additionally, we thank Tristan Rodriguez for providing the *Bmpr1a*^−/−^ ESCs used within this study. SMM was supported by a Wellcome Trust Sir Henry Wellcome postdoctoral fellowship under the supervision of AKH. Work in the Hadjantonakis lab was supported by grants from the NIH (R01HD094868, R01DK084391 and P30CA008748).

## Ethics

Animal experimentation: Animal experimentation: All mice used in this study were maintained in accordance with the guidelines of the Memorial Sloan Kettering Cancer Center (MSKCC) Institutional Animal Care and Use Committee (IACUC) under protocol number 03-12-017 (PI Hadjantonakis).

